# ZDHHC13 is a likely pseudoenzyme protein S-acyltransferase that functions via a non-canonical mechanism

**DOI:** 10.64898/2026.04.20.719575

**Authors:** Andrey A. Petropavlovskiy, Alysha M. Church, Amelia H. Doerksen, Denver A. Bakhareva, Étienne P. Sellar, Nisandi N. Herath, Shaun S. Sanders

## Abstract

S-acylation is the addition of fatty acids to cysteine residues to regulate protein function and localization. S-acylation is catalyzed by the ZDHHC (Asp–His–His–Cys) family of protein S-acyltransferases (PATs), which S-acylate protein substrates by first auto-S-acylating the catalytic cysteine of the DHHC active site followed by transfer to the substrate. ZDHHC13 and ZDHHC17 are related ankyrin repeat domain (ANK) PATs that S-acylate multiple neuronal proteins, including huntingtin (HTT), the protein mutated in Huntington disease. However, unlike ZDHHC17 and other human PATs, ZDHHC13 possesses a non-canonical DQHC active site. As the first histidine is essential for auto-S-acylation, it is unclear if ZDHHC13 is catalytically active. Our phylogenetic analysis of eukaryotic ANK-containing PATs shows that ZDHHC13 orthologues are more divergent compared to ZDHHC17. While the ZDHHC17 DHHC is highly conserved, the motif varies among ZDHHC13 orthologues, with some vertebrate lineages containing a serine in place of the catalytic cysteine. Interestingly, we found that the ZDHHC13 S-acylation is lower than that of ZDHHC17, but the ZDHHC13 catalytic cysteine is indeed S-acylated. While expression of wild type (WT) ZDHHC13 in *ZDHHC13* deficient HEK293T cells increased S-acylation of a HTT^1-588^ fragment, surprisingly, expression of catalytically dead DQHS ZDHHC13 was still able to facilitate HTT^1-588^ S-acylation equally. This suggests the ZDHHC13 catalytic cysteine is not required for S-acylation of target proteins, suggesting ZDHHC13 may coordinate another PAT. Indeed, we identified ZDHHC13 in high-molecular weight complexes. Our results indicate that ZDHHC13 is a likely pseudoenzyme that may function via a non-conventional mechanism reliant on other PATs. This work broadens our understanding of the function of this non-canonical PAT.

---

Precise subcellular protein trafficking and localization is critical for cellular function. One mechanism that regulates cellular protein trafficking is the post translational lipid modification S-acylation (palmitoylation). S-acylation is the addition of long-chain fatty acids to protein cysteine residues via a labile thioester bond. S-acylation thereby regulates the interaction of proteins with lipid membranes, altering protein transport, sorting, protein-protein interactions, and function.^1^ Due to its critical role in many cellular processes, altered S-acylation has been implicated in human disease, including neurological disorders, cancers, metabolic disorders, and immune function and infections.^1–3^

The 23 member ZDHHC family of protein S-acyltransferases (PATs) catalyze S-acylation and are characterized by their conserved Asp–His–His–Cys (DHHC) active site motif within a zinc-finger cysteine-rich domain (CRD).^1,4^ PATs are transmembrane proteins with 4-6 α-helical transmembrane domains with their DHHC motif active site located in a β-sheet facing the cytosol.^5,6^ PATs S-acylate target proteins via a two-step process facilitated by a catalytic triad of His, Asp, and Cys.^6,7^ The enzyme auto-S-acylates (simply autoacylates hereto in) on the Cys of the catalytic triad. During autoacylation, the first His of the DHHC motif acts as a Brønstead-Lowry base to deprotonate the Cys converting the thiol into a thiolate nucleophile to allow autoacylation by nucleophilic attack on the carbonyl carbon of acyl-CoA. The acyl group is then transferred to the target substrate. As such, changing the His to Ala or the Cys to Ser renders ZDHHCs catalytically dead *in vitro*.^6^

ZDHHC17 (Huntingtin-interacting protein 14 [HIP14]) and ZDHHC13 (HIP14-Like [HIP14L]) are related PATs and the only PATs that have six transmembrane domains and amino (N)-terminal ankyrin repeat domains (ANK)^8–11^ They were initially discovered as PATs for Huntingtin (HTT), the protein mutated in Huntington disease.^2^ Interestingly, the DHHC active site motif that is conserved across all the other 22 human PATs is not shared by ZDHHC13. Instead of a His at residue 454, the active site motif of ZDHHC13 has a glutamine (DQHC rather than DHHC).^9^ His to Gln is not a synonymous substitution, as these residues are far from each other structurally and biochemically.^12^ As such, Gln in the active site of ZDHHC13 is likely unable to deprotonate the Cys. Furthermore, based on the AlphaFold2 predicted structure, ZDHHC13 does not have a residue in proximity to the Cys that could substitute functionally for His in this capacity.^13^

While S-acyl transferase activity of ZDHHC17 towards synaptosome associated protein 25 (SNAP25) has been demonstrated *in vitro* with purified protein,^11,14^ activity of ZDHHC13 has only been observed indirectly from *in cellulo* or *in vivo* studies. Co-overexpression of ZDHHC13 with putative substrates, such as HTT or ClipR-59 (CLIP-170 related protein), leads to increased S-acylation while loss of ZDHHC13 results in reduced S-acylation of putative substrates.^8,14–16^ Thus, S-acylation of several proteins is clearly dependent on ZDHHC13 but how the DQHC active site motif affects ZDHHC13 S-acyltransferase activity and if it is a catalytically active enzyme remains unknown.

To address this question, we sought to determine if ZDHHC13 is catalytically active compared to a catalytically dead DQHS (C456S) variant and if changing the glutamine in ZDHHC13 active site to the canonical histidine restores and/or increases its catalytic activity.

## Results & Discussion

### Mammalian ZDHHC13 likely evolved as a pseudoenzyme

To better understand the evolution of ZDHHC13 in relation to ZDHHC17 and other eukaryotic ANK ZDHHC proteins, we collected and aligned all ANK ZDHHC sequences from a set of model organisms across eukaryotic kingdoms and generated a maximum likelihood phylogenetic tree based on this alignment (Figure S1 and S2). Using the phylogenetic tree, we then separated the ZDHHC17 and ZDHHC13 orthologues for their comparison. The CRD in ZDHHC13 orthologues in human (*Homo sapiens*), house mouse (*Mus musculus*), chicken (*Gallus gallus*), western clawed frog (*Xenopus tropicalis*), and zebrafish (*Danio rerio*) have low sequence identity with only 53% between human and zebrafish (Figure 1A). In contrast, there is high sequence identity ranging from 93-100% within the CRD of ZDHHC17 orthologues (Figure 1A). Notably, a representative cyclostome, inshore hagfish (*Eptatretus burgeri*), possesses a ZDHHC17 orthologue (A0A8C4Q4D6), but no ZDHHC13 orthologue (Figure 1A). The ZDHHC13 active site motif is DQHC in human, mouse, and frog, DQHS in chicken, and DHHS in zebrafish, none of which are predicted to be capable of catalysis (Figure 1A). In contrast, the ZDHHC17 orthologue active site motif is DHHC in all sequences. While some ANK ZDHHCs contain a DHYC active site (e.g. *Saccharomyces cerevisiae*), the ZDHHC13 orthologues branch is the only in which variation occurs at the first His or the catalytic Cys.

**Figure 1.**
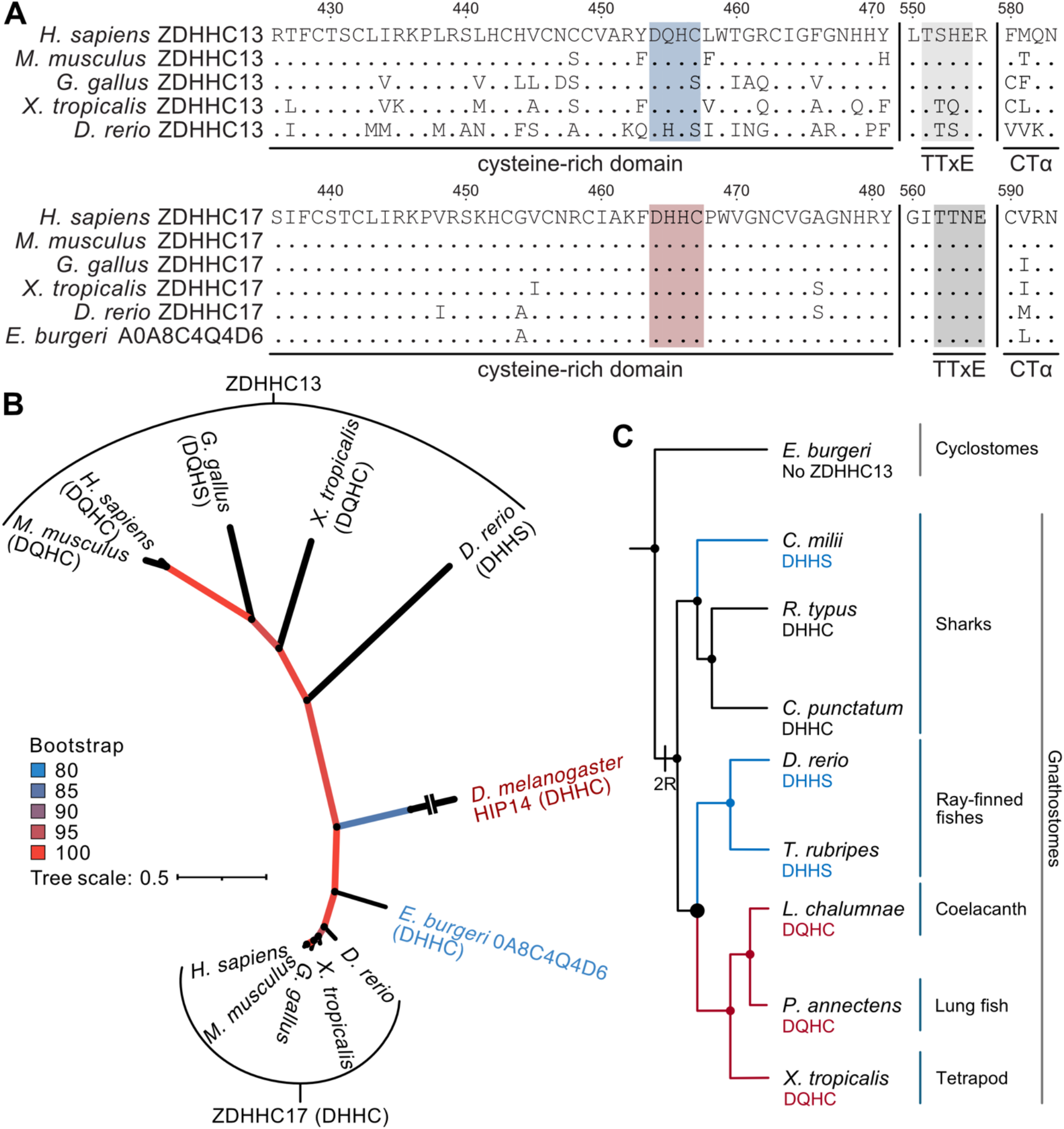
ZDHHC13 orthologues appeared early in vertebrate evolution and evolved differently compared to ZDHHC17. (A) Multiple sequence alignment of the CRD of ZDHHC13 (top) and ZDHHC17 (bottom) from vertebrate with the catalytic motif highlighted in blue (ZDHHC13) or red (ZDHHC17), the TTxE region highlighted in grey, and four residues of the C-terminal α-helix (CTα) PaCCT motif shown. (B) Pruned maximum-likelihood phylogenetic tree reconstructed using IQTree 3.0.1 showing the relationship between ZDHHC13 and ZDHHC17 orthologues. Line colour indicates bootstrap values as in the legend, active site motif sequences are in brackets, and species names are coloured as follows: vertebrates in black, cyclostomes in blue, and insects in maroon. Two dashes indicate branch were shortened for fit. (C) Cladogram of early vertebrate evolution with the sequences of catalytic motifs indicated. 2R indicates the second vertebrate whole genome duplication event in the gnathostome lineage after which ZDHHC13 orthologues appear. Species with a DHHS active site motif are indicated in blue and those with DQHC in maroon with the thick node indicating the split point between DHHS and DQHC. Cladogram adapted from Yu *et al*., 2024.^19^

While the carboxyl (C)-termini of ZDHHC enzymes are not well conserved, there is a highly conserved TTxE motif (Thr-Thr-x-Glu, where x is any amino acid). Human, mouse, and chicken ZDHHC13 have a Thr to Ser substitution in second position of TTxE motif (Figure 1A), which, in the ZDHHC20 structure, stabilizes the DHHC.^6^ The C-terminus of ZDHHCs also contain the PaCCT (palmitoyltransferase conserved C-terminus) motif with a highly conserved Asn at position 11 (N583 in human ZDHHC13).^17^ ZDHHC13 N583 is conserved across vertebrate orthologues (Figure 1A).

Based on the maximum-likelihood phylogenetic tree reconstructed from full-length alignment, while ZDHHC17 orthologues cluster closely together, ZDHHC13 orthologues are more divergent both from each other and from the ZDHHC13/ZDHHC17 common ancestor (Figure 1B), consistent with a previous metazoan ANK ZDHHC phylogeny.^18^ Vertebrate ZDHHC13 and ZDHHC17 diverged from a common ancestor close to that of *Drosophila melanogaster* HIP14. Indeed, fish ZDHHC13 is the closest relative to the common ancestor and to the ZDHHC17 clade (Figures 1B & S2). The other ANK ZDHHC proteins are divergent from both vertebrate ZDHHC17 and ZDHHC13, forming several distinct clades, however, the relationship between these clades is more difficult to establish due to low bootstrap values (Figure S2).

As we noticed substantial variation in ZDHHC13 active site sequences among fishes, we further examined early ZDHHC13 evolution by correlating the active site motif sequence with speciation events based on vertebrate protein phylogeny.^19^ Alignment of fish sequences shows that while most shark ZDHHC13 orthologues contain a DHHC motif, all ray-finned fish ZDHHC13 orthologues contain a DHHS motif, whereas coelacanth, lungfish, and tetrapod branches have a DQHC motif (Figures 1C, S1, & S3). Thus, the origins of the DQHC active site can be traced to the lobe-finned and tetrapod divergence. Based on the vertebrate cladogram and this alignment we can speculate on the origins of vertebrate ZDHHC13. As there are no ZDHHC13 orthologues in cyclostomes, but they do exists in all other fishes (Figure 1C), ZDHHC13 could have originated from an ancestral ANK ZDHHC during the last vertebrate whole genome duplication event (2R) after divergence of cyclostomes and gnathostomes.^19^ Following this duplication the catalytic function of ZDHHC13 may have become redundant, reducing the selective pressure to retain the DHHC active site, allowing the inactivating substitutions to occur. This may have also allowed ZDHHC13 to subspecialize and gain new functions.

Given the His454 to Gln substitution that is likely to render ZDHHC13 incapable of autoacylation, and higher variation in the ZDHHC13 CRD and DHHC, TTxE, and PaCCT motifs, we suggest that ZDHHC13 may be a pseudoenzyme. Pseudoenzymes are catalytically deficient proteins that share a common ancestor and structure with their catalytically active counterparts.^20^ They often have higher sequence variation than their active cognates^21^, and originate from a catalytically active ancestor through gene duplication and subsequent diversification.^20^ The fact that a DQHC motif is a unique feature of vertebrate ZDHHC13 orthologues also supports the pseudoenzyme designation. While the second His is more variable across ZDHHC13 and ZDHHC17 orthologues, with some branches having Tyr in its place (Figures S1 & S2), His in the first position is highly conserved across all branches. Consistent with this, computational prediction^22^ of enzymatic activity shows the highest K_m_ for ZDHHC13 compared to that of ZDHHC17 and two highly active ZDHHC enzymes, ZDHHC3 and ZDHHC20^5,6,23^ (Figure 2A).

**Figure 2.**
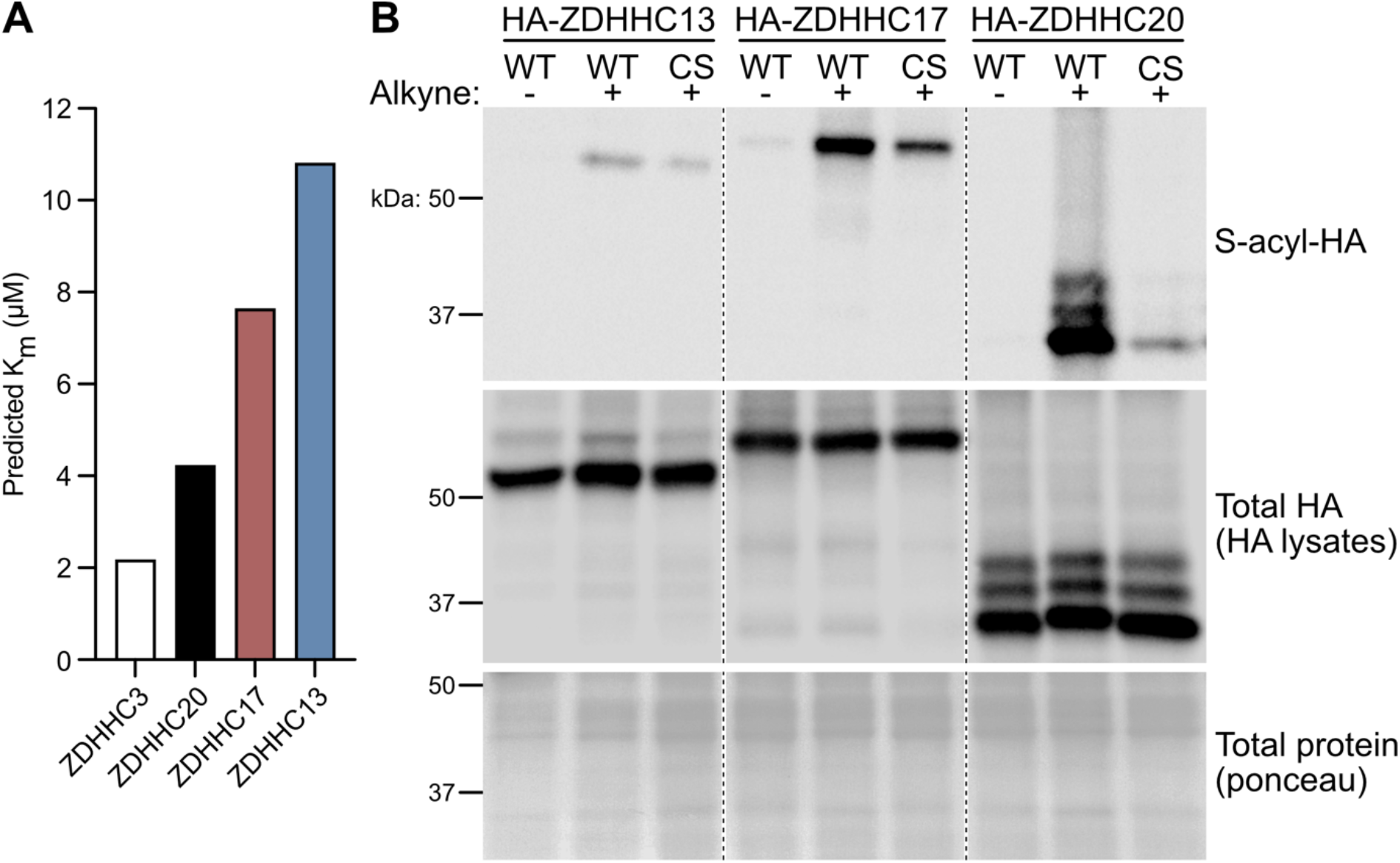
ZDHHC13 is less S-acylated than ZDHHC17 or ZDHHC20 but all three are S-acylated on the catalytic Cys. (A) K_m_ values were predicted for ZDHHC3, ZDHHC13, ZDHHC17, and ZDHHC20 using CatPred^*22*^ with deprotonated palmitoyl-CoA as the substrate. (B) HEK293T cells were transfected to express HA-tagged ZDHHC13 WT or CS, ZDHHC17 WT or CS, or ZDHHC20 WT or CS and labelled with alkyne-palmitate or palmitate (negative control) for 1 hour. S-acylated proteins were purified from lysates using click chemistry and detected by immunoblot using an HA antibody (top panel). Total HA levels in parent lysates were also determined (middle panel) as well as total protein using ponceau stain (bottom panel). Composite images are from the same immunoblot image and cuts are indicated by dotted lines. Blots are representative of 3 independent experiments.

### The ZDHHC13 catalytic Cys is S-acylated

Thus, we next sought to experimentally determine if ZDHHC13 is catalytically active or not, as we predict based on our bioinformatic analyses. First, we compared S-acylation levels of wild type (WT) and catalytically dead (Cys to Ser, CS) variants of ZDHHC13 (DQHS; C456S) to that of ZDHHC17 (DHHS; C467S) and ZDHHC20 (DHHS; C156S) using bioorthogonal labeling with alkyne-palmitate and click chemistry in human embryonic kidney 293T cells (HEK293T) (Figure 2B). The S-acylation level of WT ZDHHCs compared to CS variants is thus indicative of catalytic cysteine S-acylation. ZDHHC20 displayed the highest S-acylation levels of the three enzymes and ZDHHC13 the lowest (Figure 2B), consistent with our computational prediction of enzyme activity (Figure 2A). Furthermore, all three CS variants are still S-acylated to some extent, with ZDHHC17 CS S-acylated the strongest, suggesting modification of other Cys residues (Figure 2B). While ZDHHC13 is S-acylated at the catalytic Cys, this may not be due to autoacylation. Given that other cysteine residues on ZDHHC13 are S-acylated, it is also possible that the ZDHHC13 catalytic Cys is S-acylated by another enzyme in a PAT cascade. Indeed, a PAT cascade involving regulation of ZDHHC6 by ZDHHC16-mediated S-acylation has been demonstrated.^24^ Thus, this type of mechanism may account for ZDHHC13 S-acylation at the DQHC Cys residue.

### Changing the ZDHHC13 DQHC Gln to a canonical His does not increase its S-acylation

We next sought to determine whether S-acylation of the catalytic cysteine can be enhanced by changing the Gln to a canonical His. To this end, we compared S-acylation levels of WT, CS, and QH (DHHC; Q454H) variants of ZDHHC13 using bioorthogonal labeling and click chemistry (Figure 3A). The ZDHHC13 CS variant was 42% less S-acylated than WT (Figure 3A&B). Surprisingly, S-acylation of the ZDHHC13 QH variant was reduced to a similar level, by 46% (Figure 3A&B). We compared these results to WT and CS ZDHHC17 as well as a variant with the first His changed to a Gln (DQHC; H465Q). ZDHHC17 CS S-acylation was reduced by 67% and that of the HQ variant by 49%, consistent with the canonical catalytic role of the first His of the DHHC motif expected for ZDHHC17 (Figure 3C&D). The failure of the QH substitution to enhance S-acylation of the catalytic Cys suggests that other parts of the protein are unable to support autoacylation activity, perhaps the lack of a canonical TTxE (TSHE in human and mouse ZDHHC13) or other variations in the CRD. Alternatively, it is possible that ZDHHC13 operates using a completely different mechanism that requires the Gln for S-acylation of Cys.

**Figure 3.**
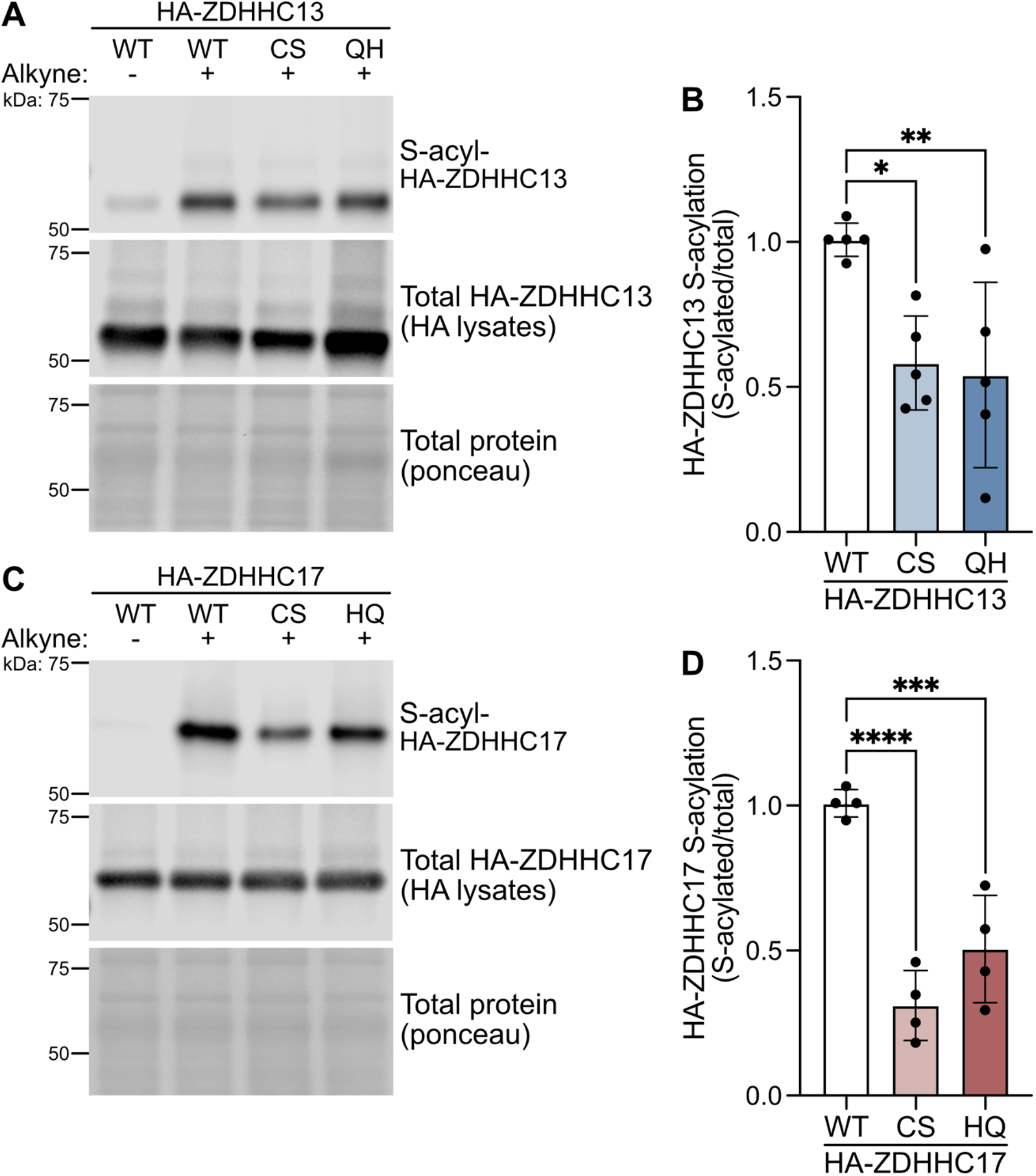
Gln/His and Cys active site residues of ZDHHC13 and ZDHHC17 are required for their S-acylation. (A) HEK293T cells expressing HA-ZDHHC13 WT, CS, or QH were labelled with alkyne-palmitate or palmitate (negative control) for one hour. S-acylated proteins were purified from lysates using click chemistry and detected by immunoblot using an HA antibody (top panel). Total HA levels in parent lysates were also determined (middle panel) as well as total protein using ponceau stain (bottom panel). (B) Quantified data from A, showing S-acyl/total levels of HA-ZDHHC13 normalized to WT (one-way ANOVA p=0.0075, F(2,12)=7.57, N=5, Dunnett’s multiple comparison WT vs CS * p=0.014, WT vs QH ** p=0.0079). (C) HEK293T cells expressing HA-ZDHHC17 WT, CS, or HQ were labelled with alkyne-palmitate or palmitate (negative control) for one hour. S-acylated proteins were purified from lysates using click chemistry and detected by immunoblot using an HA antibody (top panel). Total HA levels in parent lysates were also determined (middle panel) as well as total protein using ponceau stain (bottom panel). (D) Quantified data from C, showing S-acyl/total levels of HA-ZDHHC17 normalized to WT (one-way ANOVA p<0.0001, F(2,9)=30.41, N=4, Dunnett’s multiple comparison WT vs CS **** p<0.0001, WT vs HQ *** p=0.00080).

### S-acylation of ZDHHC13 substrates is not dependent on its catalytic activity

Given our data that ZDHHC13 is likely unable to autoacylate while being S-acylated on the DQHC Cys, we next sought to determine if ZDHHC13 can transfer acyl groups to a target substrate protein. To do so, we first evaluated S-acylation of peptidyl arginine deaminase type III (PADi3), a proposed ZDHHC13 substrate. In 2020, Chen *et al*. reported decreased S-acylation of PADi3 in *Zdhhc13* Arg452stop homozygous mice and one replicate showing increased S-acylation in HEK293T cells when co-expressed with WT mouse ZDHHC13.^25^ However, using bioorthogonal labeling and click chemistry or the acyl biotin exchange (ABE) assay in HEK293T cells, we could not replicate the reported increase in PADi3 S-acylation (Figure S4 and data not shown). As HTT is a well-established substrate of both ZDHHC13 and ZDHHC17^14^, we compared S-acylation of an eYFP (yellow fluorescent protein)-tagged, N-terminal 1-588 amino acid HTT fragment with co-expression of WT, CS, or QH ZDHHC13 using the ABE assay. HTT S-acylation is robustly enhanced ∼4-fold in cells when co-expressed with WT ZDHHC13 (Figure 4A&B). Surprisingly, both the CS and QH ZDHHC13 variants also enhanced HTT^1-588^-YFP S-acylation to a similar level (Figure 4A&B). This suggests that while HTT S-acylation is dependent on ZDHHC13 levels, ZDHHC13 facilitates S-acylation through a non-canonical mechanism not requiring the catalytic Cys. It is possible that ZDHHC13 directly catalyzes S-acylation without involving the DQHC or that ZDHHC13 coordinates another PAT to facilitate S-acylation of substrate proteins. Indeed, this theory was proposed by the Chamberlain group based on their data that neither WT nor QH ZDHHC13 increased S-acylation of the ZDHHC17 substrate SNAP25^26^, despite reduced S-acylation of SNAP25 in the brain of *Zdhhc13* knockout mice.^8^

**Figure 4.**
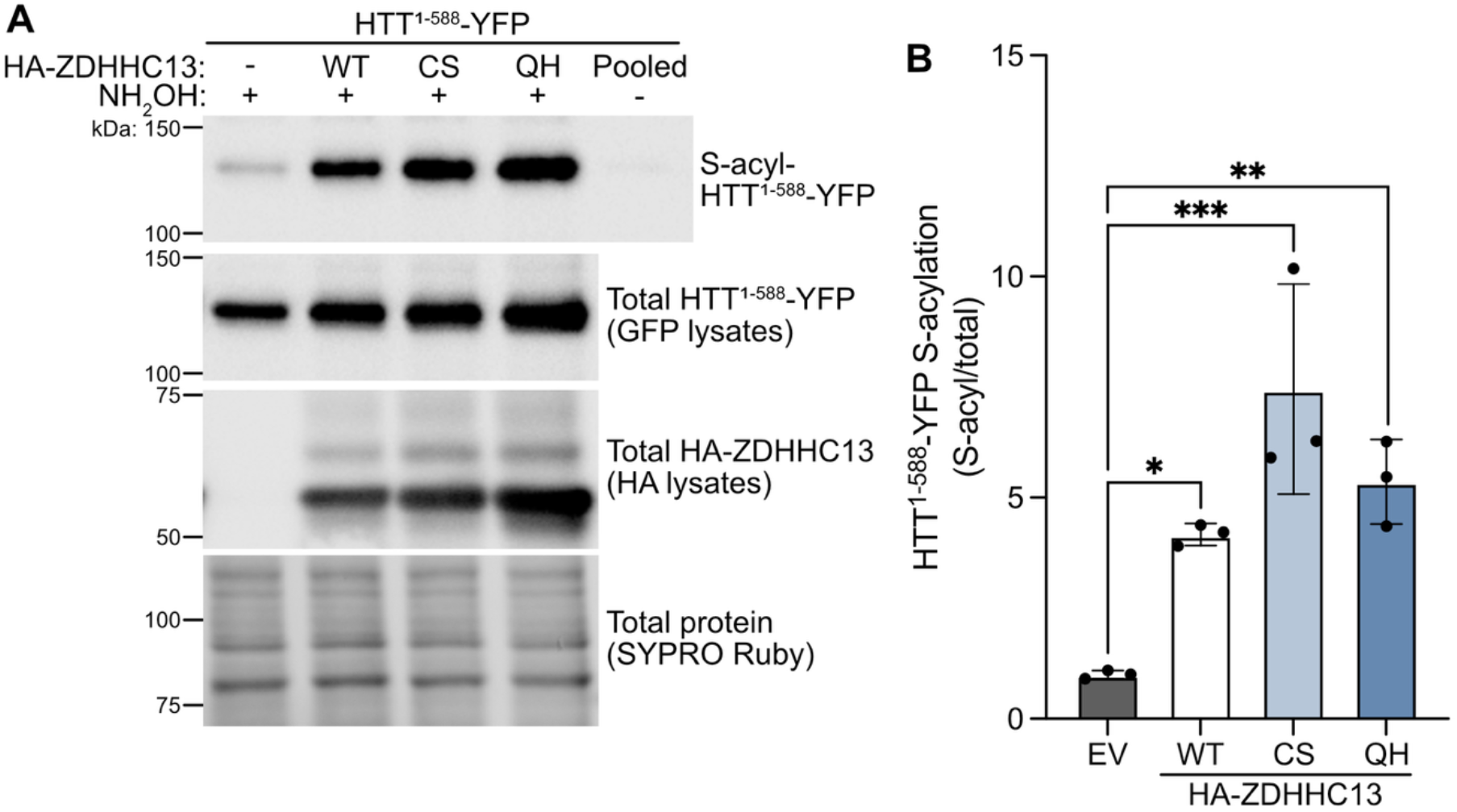
ZDHHC13 WT, CS, and QH variants S-acylate HTT fragment equally. (A) S-acylated proteins were purified from HEK293T cells expressing HTT^1-588^-YFP alone (empty vector [EV] or -) or with HA-ZDHHC13 WT, CS, or QH by ABE assay and detected by immunoblot using an antibody against GFP (top panel). A parallel pooled sample processed in the absence of the key ABE reagent hydroxylamine (NH_2_OH) confirm the specificity of the ABE assay. Total GFP and HA levels in parent lysates were also determined (middle panels) as well as total protein using SYPRO Ruby stain (bottom panel). (B) Quantified data from A, showing S-acyl/total levels of HTT^1-588^-YFP normalized to EV (one-way ANOVA p=0.0018, F(3,8)=13.21, N=3, Dunnett’s multiple comparison EV vs WT * p=0.041, EV vs CS *** p=0.0007, EV vs QH ** p=0.0081).

It may be that endogenous ZDHHC13 facilitates S-acylation of HTT^1-588^-YFP in Figure 4, thus masking differences in S-acylation by overexpressed WT, CS, or QH ZDHHC13. To address this, we generated *ZDHHC13* knockout cells using a CRISPR-Cas9 dual guide RNA (gRNA) strategy. We verified loss of ZDHHC13 in the knockout cells by immunoblot (Figure 5A&B). We then assessed HTT^1-588^-YFP S-acylation with co-expression of WT, CS, and QH ZDHHC13 in NT (non-targeting gRNA) and *ZDHHC13* knockout cells using the ABE assay. As in the WT HEK293T cells, all variants of ZDHHC13 were able to enhance HTT^1-588^-YFP S-acylation to the same levels in *ZDHHC13* knockout and NT cells (Figure 5C&D). Thus, our findings in Figure 4 were not due to S-acylation by endogenous ZDHHC13, and ZDHHC13 likely coordinates another PAT to facilitate S-acylation of substrates. The most logical candidate PAT whose activity is facilitated by overexpressed ZDHHC13 is ZDHHC17, the other ANK PAT. To test this hypothesis, we generated *ZDHHC17* knockout cells using the same CRISPR-Cas9 approach as above. We verified loss of ZDHHC17 in the knockout cells (Figure 6A&B) and assessed S-acylation of HTT^1-588^-YFP with co-expression of WT, CS, and QH ZDHHC13 in NT and *ZDHHC17* knockout cells using the ABE assay. Similar to findings in Figure 5, all variants of ZDHHC13 enhanced HTT^1-588^-YFP S-acylation to the same levels in both NT and ZDHHC17 knockout cell lines (Figure 6A&B). This suggests that ZDHHC17 is not necessary for S-acylation of HTT^1-588^-YFP by ZDHHC13 and that ZDHHC13 is able to coordinate another, non-ANK ZDHHC.

**Figure 5.**
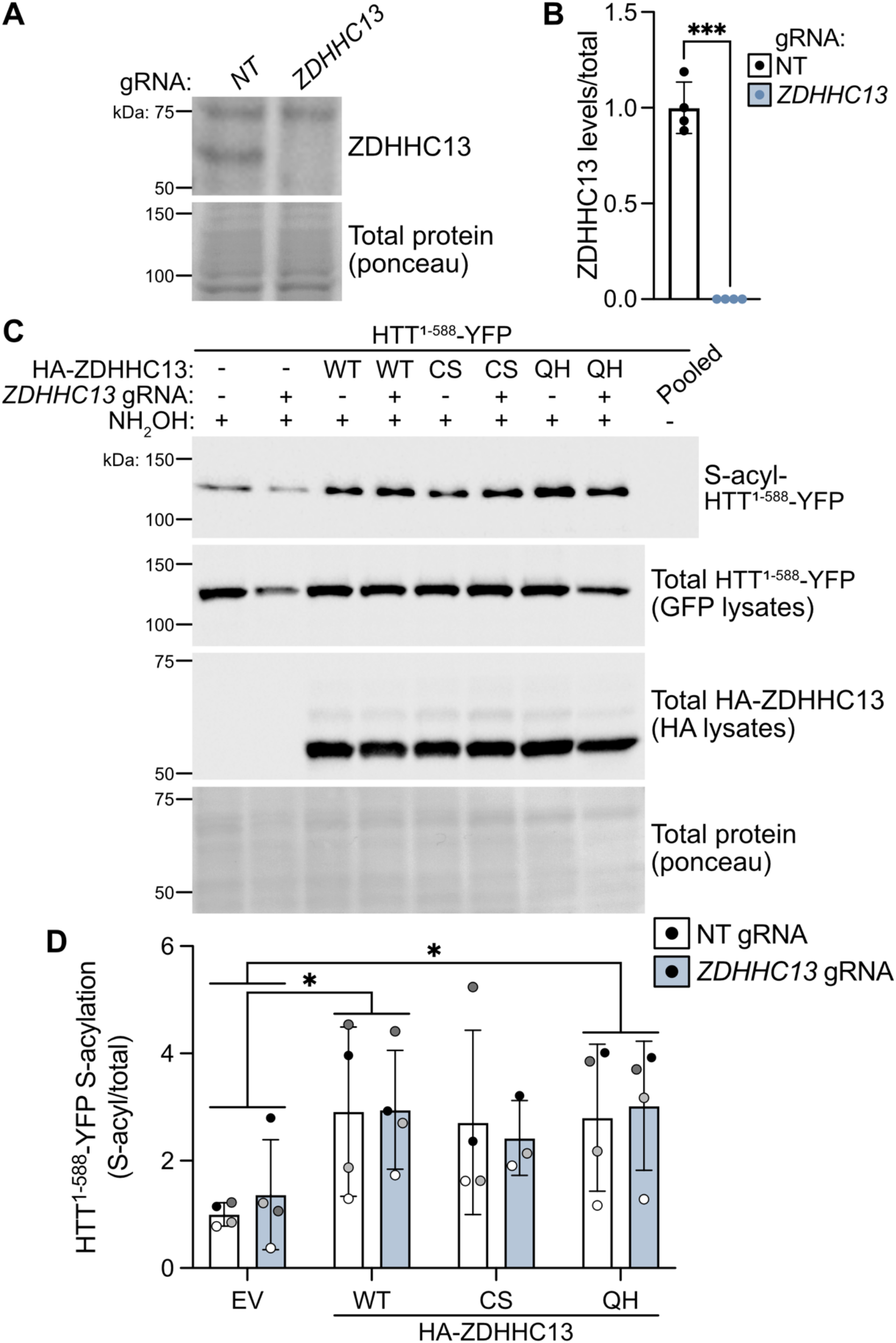
Endogenous ZDHHC13 does not account for the ability of HA-ZDHHC13 CS and QH to enhance S-acylation of HTT^1-588^-YFP. (A) ZDHHC13 levels in lysates from NT (non-targeting gRNA) and *ZDHHC13* knockout HEK293T cells were determined by immunoblot using an antibody against ZDHHC13 (top panel). Total protein levels using ponceau stain were determined (bottom panel). (B) Quantified data from A, showing ZDHHC13 levels relative to total protein normalized to NT cells (Welch’s unpaired t test p=0.0007, N=4). (C) S-acylated proteins were purified from *ZDHHC13* knockout or NT HEK293T cells expressing HTT^1-588^-YFP alone or with HA-ZDHHC13 WT, CS, or QH by ABE assay and detected by immunoblot using an antibody against GFP (top panel). Total GFP and HA levels in parent lysates were also determined (middle panels) as well as total protein (ponceau, bottom panel). (D) Quantified data from C, showing S-acyl/total levels of HTT^1-588^-YFP normalized to EV (Two-Way ANOVA HA-ZDHHC13 variant p=0.027 [F(3, 23)=3.68], cell line not significant (ns), interaction ns, N=3-4, Dunnett’s multiple comparison EV vs WT * p=0.023, EV vs QH * p=0.025).

**Figure 6.**
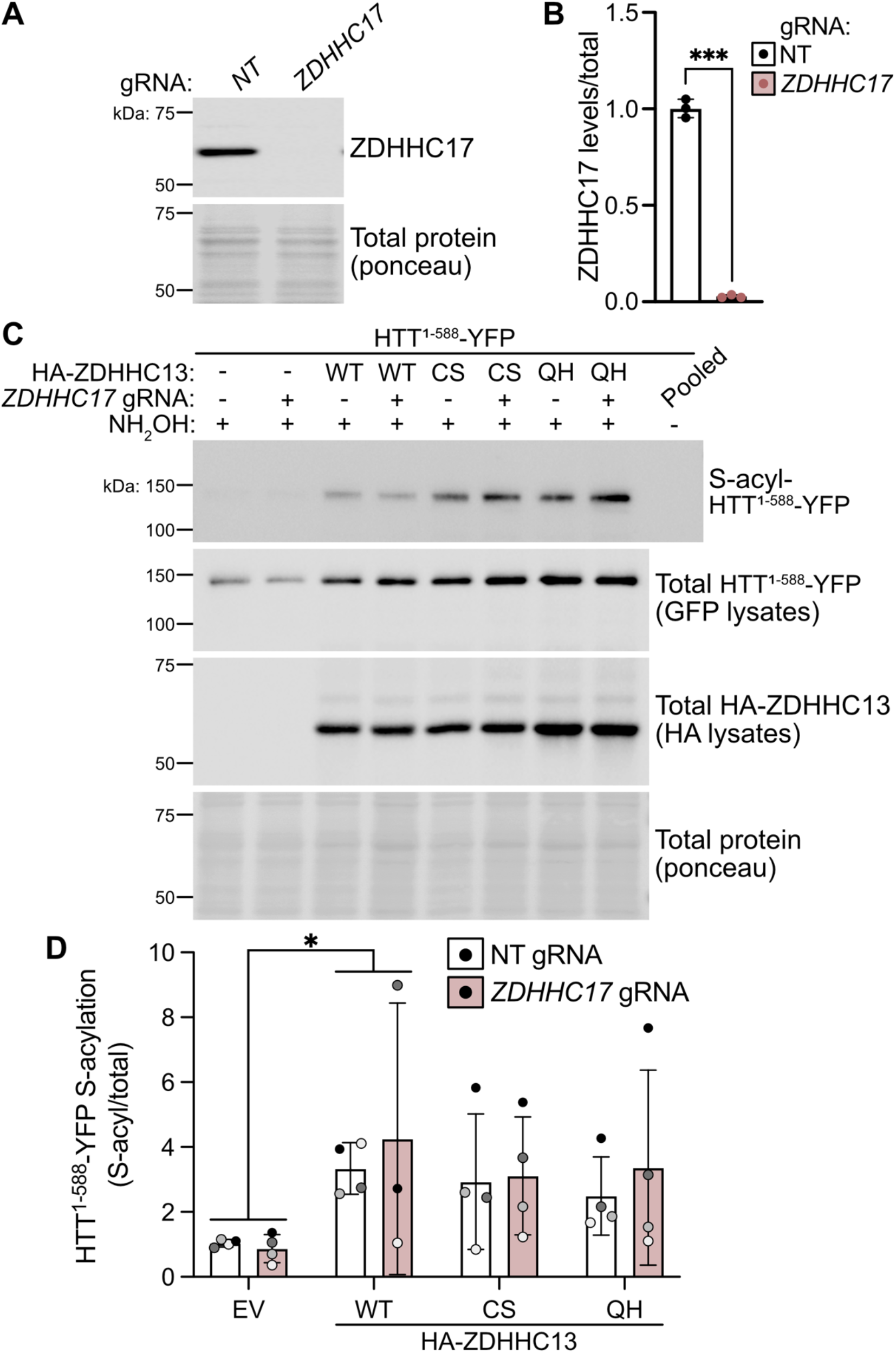
Endogenous ZDHHC17 does not account for the ability of HA-ZDHHC13 CS and QH to enhance HTT^1-588^-YFP S-acylation. (A) ZDHHC17 levels in lysates from NT and *ZDHHC17* knockout HEK293T cells were determined by immunoblot using an antibody against ZDHHC17 (top panel). Total protein using ponceau stain was determined (bottom panel). (B) Quantified data from A, showing ZDHHC17 levels relative to total protein normalized to NT cells (Welch’s unpaired t test p=0.0006, N=3). (C) S-acylated proteins were purified from *ZDHHC17* knockout or NT HEK293T cells expressing HTT^1-588^-YFP alone or with HA-ZDHHC13 WT, CS, or QH by ABE assay and detected by immunoblot using an antibody against GFP (top panel). Total GFP and HA levels in parent lysates were also determined (middle panels) as well as total protein (ponceau, bottom panel). (D) Quantified data from C, showing S-acyl/total levels of HTT^1-588^-YFP normalized to EV (Two-Way ANOVA HA-ZDHHC13 variant ns, cell line ns, interaction ns, N=3-4, Dunnett’s multiple comparison EV vs WT * p=0.032).

### ZDHHC13 likely scaffolds a multimeric protein complex

If ZDHHC13 can facilitate other ZDHHC enzymes, we might expect that it would be doing so as a subunit of a multimeric complex. To determine if ZDHHC13 is present in high molecular weight complexes we used blue native PAGE (polyacrylamide gel electrophoresis) to resolve native protein complexes. Indeed, all variants of ZDHHC13 are present in a ∼200 kDa complex (Figure 7A&B). The size of this complex is consistent with more subunits than simple ZDHHC13/ZDHHC13 or ZDHHC13/ZDHHC17 dimers as HA-ZDHHC13 is ∼60 kDa and HA-ZDHHC17 ∼65 kDa. While it is possible that ZDHHC17 is present in the complex with ZDHHC13, the size of the complex suggests that another non-ANK PAT and/or other proteins are also present, consistent with our finding that ZDHHC17 is not required for ZDHHC13 activity (Figure 6). Indeed, there may be other proteins present in the complex including substrates and/or adaptor or accessory proteins. ZDHHC2 and ZDHHC3 have been shown to form dimers, negatively correlated with activity levels where inactive enzyme was correlated with dimer formation.^27,28^ Similar results have been reported for the yeast ZDHHC enzyme Swf1.^29^ ZDHHC17 also interacted with itself, but not ZDHHC13, in a yeast 2-hybrid screen, but this was not verified using low-throughput methods or correlated with activity levels.^30^

**Figure 7.**
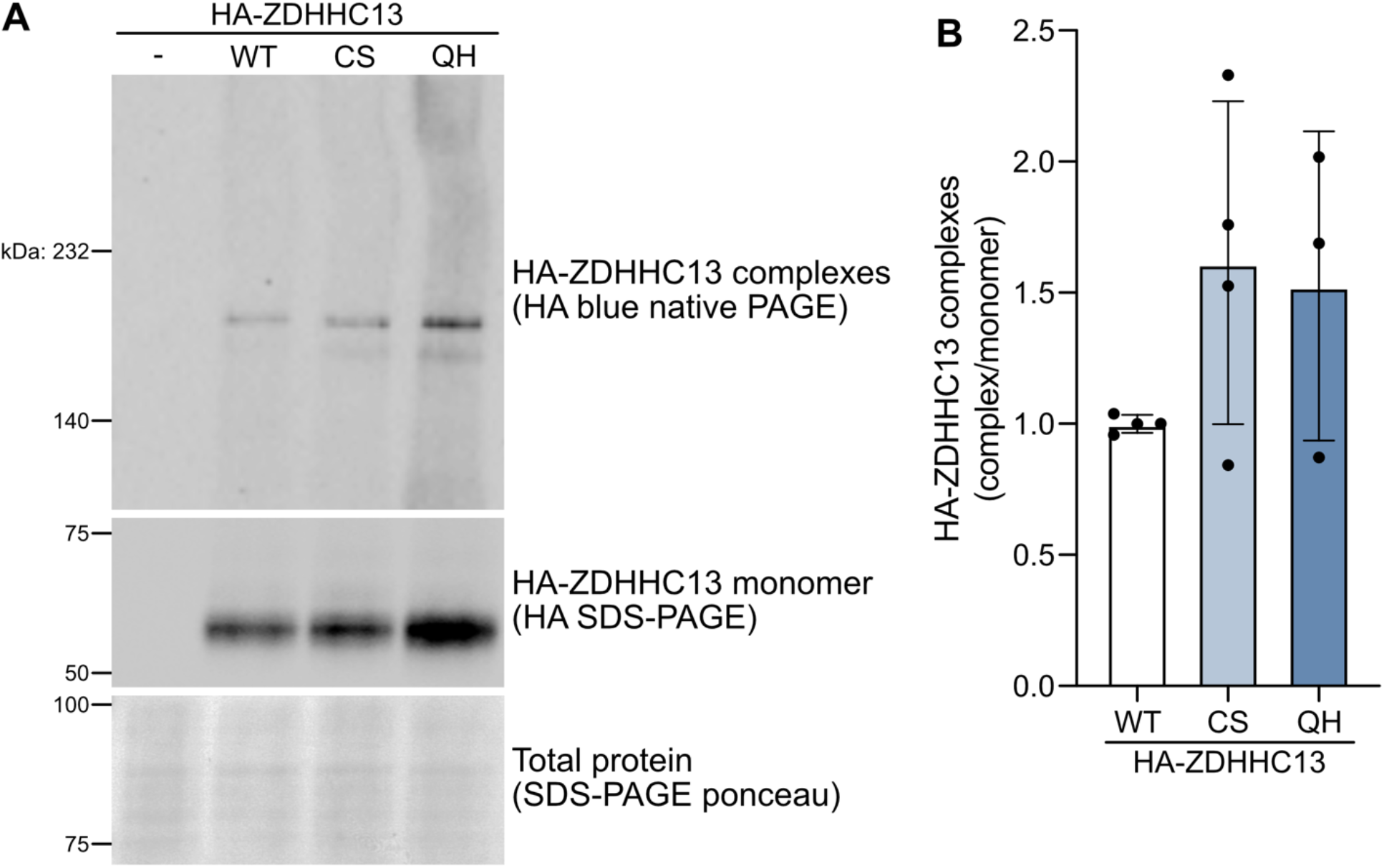
ZDHHC13 is present in high molecular weight complexes. (A) Crude membranes were isolated from HEK293T cells expressing HA-ZDHHC13 WT, CS, or QH and subjected to blue native PAGE (top panel) and denaturing SDS-PAGE (middle panel) followed by immunoblot using an antibody against HA. The ‘-’ condition was transfected with EV. Total protein using ponceau stain (bottom panel) was determined on the SDS-PAGE blot. (B) Quantified data from A, showing HA-ZDHHC13 levels in complexes relative to monomer levels (one-way ANOVA ns, N=2-3).

Here we provide the first substantial evolutionary and biochemical evidence that ZDHHC13 is catalytically deficient and a likely pseudoenzyme. Importantly, this is also the first time that ZDHHC13 has been shown to be in a complex. Consistent with other ZDHHCs, we see a non-significant trend to more ZDHHC13 in high molecular weight complexes with the CS and QH variants. However, we are proposing a different mechanism for ZDHHC13 in oligomeric complexes where ZDHHC13 recruits another catalytically active PAT or PATs into the complex to coordinate S-acylation of protein substrates. Perhaps ZDHHC13 acts as a local reserve of acyl groups for the active enzyme in the complex or aids in recruiting substrates. An interesting area of future work would involve identification of the other members of the complex, allowing further mechanistic studies exploring the role of ZDHHC13 in coordinating S-acylation.

Our work provides important insights into the regulation of HTT S-acylation. Indeed, enhancing S-acylation of HTT in the context of Huntington disease is protective in cell culture and mouse models^31,32^ so it is of therapeutic interest.^2^ The finding that ZDHHC13 is unlikely to directly S-acylate HTT, and instead could function as a scaffold to facilitate S-acylation by other catalytically-active ZDHHCs is important and may provide additional therapeutic targets.

## Methods

All DNA, gRNA sequence, reagent, chemical, antibody and buffer information can be found in the Key Resources Table in the Supplemental Information.

### Sequence Bioinformatics

The protein sequences of ANK ZDHHCs from common model organisms across eukaryotic kingdoms were identified using UniProt, NCBI Protein, or organism-specific databases (i.e. TAIR, Alliance of Genome Resources, TriTrypDB)^33–35^ and retrieved from UniProt (SwissProt/TrEMBL) or NCBI Protein.^36,37^ All ANK ZDHHCs identified for each specie were included in the analysis. Sequences were aligned using the T-coffee default algorithm (T-coffee, Muscle, and ClustalW; tcoffee.crg.eu)^38^ and visualized using the TexShade package (https://ctan.org/pkg/texshade).^39^ Maximum likelihood phylogenetic tree was reconstructed based on full-length alignment using the IQ-TREE package (v. 3.0.1).^40^ Prior to tree reconstruction, best substitution model was determined by ModelFinder.^41^ Bootstrap branch support values were calculated using the UFBoot algorithm, with 3000 replications and correction for severe model violation.^42^ The consensus tree was visualized using the Interactive Tree of Life (iTOL) website.^43^ CatPred implemented on TamarindBio server was used for prediction of ZDHHC K_m_ value with de-protonated palmitoyl-CoA (ChEBI:57379) as the substrate.^22^

### Molecular biology

Mutations were introduced in the cDNA sequences of pFEW-HA-*Zdhhc13*, pFEW-HA-*Zdhhc17*, and pFEW-HA-*Zdhhc20* using the Q5 site-directed mutagenesis kit using primers designed using with the NEBaseChanger Tool (https://nebasechanger.neb.com/) leading to the following amino acid changes in the DQHC/DHHC motif: ZDHHC13 Cys 456 to Ser (C456S; DQHS) and Gln 454 to His (Q454H; DHHC), ZDHHC17 Cys 467 to Ser (C467S; DHHS) and His 465 to Gln (H465Q; DQHC), and ZDHHC20 Cys 156 to Ser (C156S; DHHS). All constructs were verified by Sanger or nanopore sequencing.

### HEK293T cell culture and transfection

HEK293T cells were cultured in complete standard Dulbecco’s Modified Eagle Medium (DMEM), 10% fetal bovine serum, 1% penicillin-streptomycin, and 1% L-glutamine and maintained at 37°C in 5% CO_2_. Cells were seeded on 6-cm plates and 24-hours later transfected using CaPO_4_ as previously described.^44^ Briefly, 3.5-7 μg of DNA (3.5 μg for single transfections or 3.5 μg:0.7 μg for HA-ZDHHC13:HTT^1-588^-YFP or HA-ZDHHC13:myc-PADi3 double transfections) was diluted in 250 μL 244 mM CaCl_2_ and then combined dropwise with 250 μL 2x HEPES-Buffered Saline (HBS) with constant mixing to generate CaPO_4_-DNA precipitate. The precipitate was added dropwise to the cells, which were harvested for ABE assay, blue native PAGE, or subjected to bioorthogonal labeling (described below) 16-24 hours later.

### Gene knockout in HEK293T cells

*ZDHHC13* or *ZDHHC17* knockout HEK293T cells were generated as previously described for other genes.^45^ Cells were seeded in a 6-well plate at 62,000 cells per well, transfected the following day using CaPO_4_, as described above, with 1.2 μg of Lenti-Cas9-BLAST and 0.8 μg of pCLIP-DUAL-sgRNA plasmid. 48 hours post-transfection cells were selected using 2 μg/mL puromycin until an untransfected well displayed complete cell death. Cells were recovered into complete DMEM and maintained as a polyclonal population.

### Bioorthogonal labeling and click chemistry

20 hours following transfection, cells were subjected to bioorthogonal labeling as previously described in our protocol paper.^46^ Cells were placed in delipidated media for one hour prior to metabolic labelling with 100 μM saponified alkynyl-palmitate or non-alkyne palmitate (negative control) for one hour. Cells were lysed in modified radio-immunoprecipitation assay (RIPA) buffer with 4% Roche cOmplete EDTA (ethylenediaminetetraacetic acid)-free protease inhibitor cocktail (PIC) and protein was quantified using the Bio-Rad *DC* (Detergent Compatible) protein assay. 250 μg of lysate was mixed with 100 μM of biotin azide plus, 0.1 mM of TBTA (tris(benzyltriazolylmethyl)amine), 1mM CuSO_4_, and 1 mM sodium L-ascorbate and the reaction was carried out for one hour at room temperature prior to stopping with 10mM of EDTA pH 8. Protein was precipitated by adding ice cold acetone to a final concentration of 80%, dissolved in 2% SDS (sodium dodecyl sulfate) buffer with PIC, and diluted one in twenty in dilution buffer. Biotinylated proteins were purified using NeutrAvidin beads for three hours at 4°C prior to washing three times with dilution buffer with 0.5 M NaCl and once with dilution buffer without NaCl. S-acylated proteins were eluted using hydroxylamine (NH_2_OH) elution buffer with PIC, denatured in Laemmli sample loading buffer with 1% β-mercaptoethanol (BME) at 95°C for 5 minutes, and subjected to SDS-PAGE and immunoblotting described below.

### ABE assay

16-20 hours after transfection cells were harvested for ABE assay as previously described.^47,48^ Cells were lysed in 2% SDS buffer with 20 mM S-methyl methanethiosulfonate (MMTS) and PIC. Lysates were sonicated 2×6 seconds at 25% power and incubated 20 minutes at 50°C to block free cysteines. Protein was acetone precipitated as described above and dissolved in 4% SDS buffer with PIC. Protein was quantified using the *DC* protein assay and 300-700 μg of protein was incubated for one hour at room temperature with 0.7 M neutral hydroxylamine and 1 mM biotin-HPDP to cleave thioester bonds and biotinylate newly revealed cysteines. A pooled sample was incubated without hydroxylamine (negative control, -NH_2_OH). The protein was acetone precipitated, dissolved in 2% SDS buffer, and biotinylated proteins were affinity purified as described above. S-acylated proteins were eluted using 1% BME in elution buffer for 10 minutes at 37°C. Eluates were combined with Laemmli sample loading buffer with 1% BME and denatured at 95°C for 5 minutes and subjected to SDS-PAGE and immunoblotting described below.

### SDS-PAGE and immunoblotting

Samples were run on SDS-PAGE gels and transferred to nitrocellulose membranes. Membranes were stained for total protein using ponceau or SYPRO Ruby stain prior to blocking in 5% skim milk/Tris-buffered saline (TBS) for one hour followed by immunoblotting with the indicated primary antibodies in 1% BSA/TBS overnight at 4°C. Membranes were washed and probed with HRP-conjugated secondary antibodies in 5% skim milk/TBS for one hour at room temperature, washed, and developed with Clarity Western ECL substrate or SuperSignal West Femto Substrate. Blots were imaged using the ChemiDoc XRS+ (Bio-Rad) with Image Lab software (Bio-Rad) and quantified using Image Studio Software version 6 (Li-COR Biotech, Lincoln, NE, USA).

### Blue native PAGE

Cells were harvested in PBS and pellets frozen at -20°C. Crude membrane fractions were isolated as previously described.^49^ Cells were resuspended and incubated for 15 minutes on ice in 400 μL ice cold hypotonic buffer. 100 μL of ice cold homogenization buffer with PIC was added and cells triturated with a 26G needle 25 times, centrifuged at 1000 x g for 5 minutes at 4°C, and the supernatant ultracentrifuged at 150,000 x g for 20 minutes at 4°C to isolate the crude membrane fraction. The membrane pellet was resuspended in 150 μL of blue native PAGE lysis buffer with PIC, incubated on ice for 20 minutes, collected, and incubated at 4°C for 30 minutes with rotation. The solubilized crude membrane fraction was centrifuged at 1000 x g for 5 minutes at 4°C and the supernatant retained. Protein was quantified using the *DC* protein assay and 3-5 μg of protein was diluted in blue native lysis buffer with PIC and 0.25% Coomassie G-250. Samples were run on 4-15% SDS-free gels with cold cathode buffer in the cathode chamber and cold anode buffer in the outer chamber. The gel was destained using 20% methanol, transferred to nitrocellulose, and immunoblotted as described above. Paired samples were spiked with 1% SDS and denatured at 95°C for 5 minutes and run on SDS-PAGE as described above to determine the levels of protein monomers in the same samples.

### Statistical analysis

All data was analyzed using GraphPad Prism 10 software. All graphs show mean with standard deviation. Statistical tests used are indicated in the figure legends. Cells from separate passage numbers were considered biological replicates.

## Supporting Information

Data figures including S1 (multiple sequence alignment of CRD of ZDHHC13 and ZDHHC17 eukaryotic orthologues), S2 (phylogenetic tree of ZDHHC13 and ZDHHC17 eukaryotic orthologues), S3 (multiple sequence alignment of CRD of ZDHHC13 and ZDHHC17 fish orthologues), and S4 (ZDHHC13 does not S-acylate PADi3) as well as a key resources table containing additional methodological information including DNA, gRNA sequence, reagent, chemical, antibody and buffer information (PDF).

## Acknowledgments

Thank you to Drs. Dale Martin (University of Waterloo) and Anirban Banerjee (NIH) as well as other members of the Sanders’ Lab who assisted with helpful discussions during the preparation of this manuscript. We acknowledge that the University of Guelph resides on the Dish with One Spoon territory of the Mississaugas of the Credit of the Hodinöhsö:ni and Anishinaabe peoples, and that this land is home to many past, present, and future First Nations, Inuit, and Métis peoples.

