## Supplementary information for "ZDHHC13 is a likely pseudoenzyme protein S-acyltransferase that functions via a non-canonical mechanism"

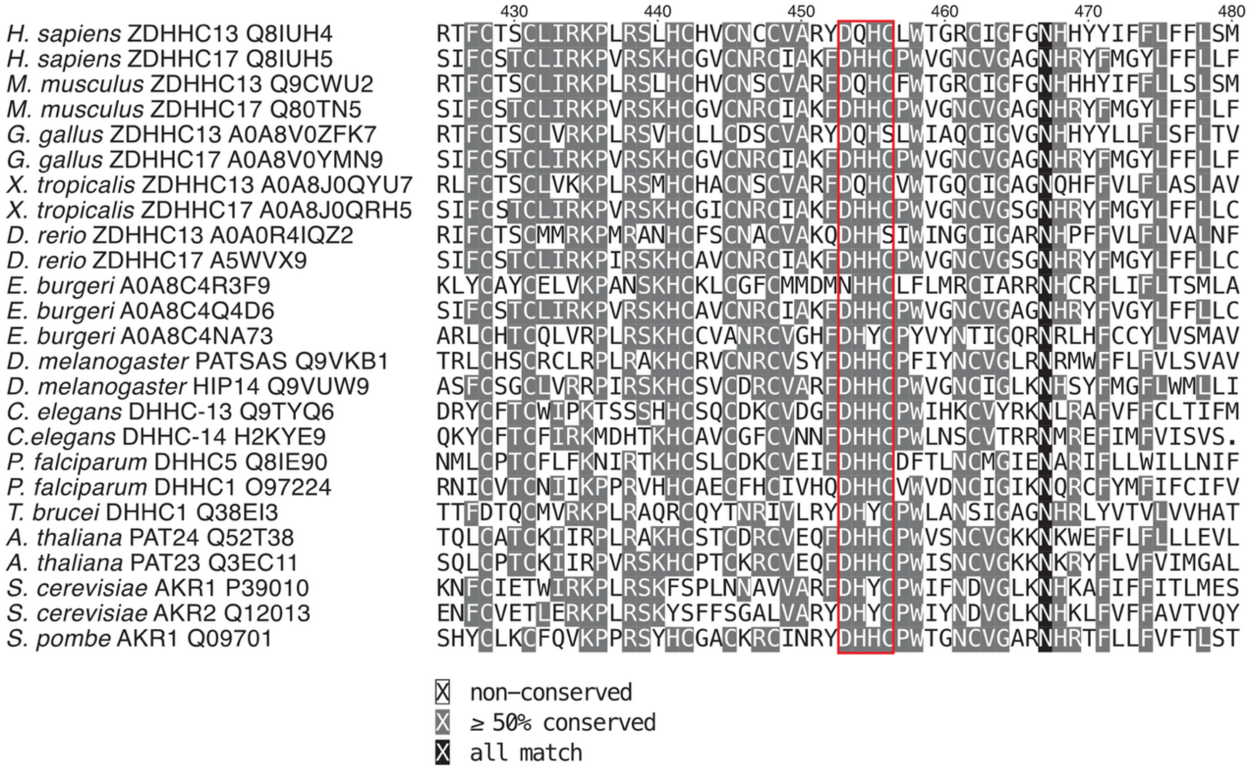

**Figure S1**. Multiple sequence alignment of the CRD of ZDHHC13 and ZDHHC17 from eukaryotic orthologues with the catalytic motif highlighted with a red box. The alphanumeric codes that follow the specie and gene names are UniProtKB accession numbers.

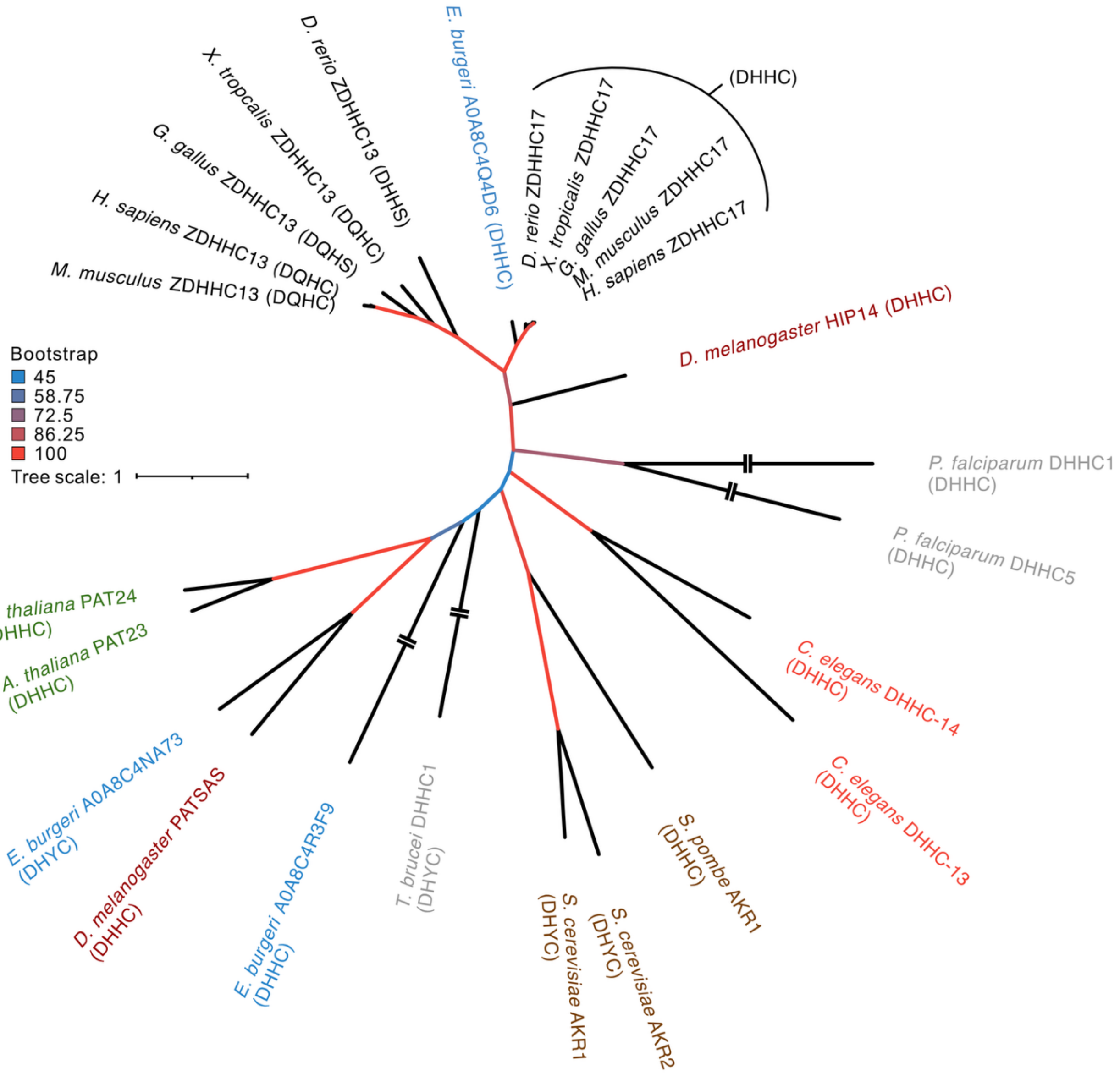


**Figure S2**. Maximum-likelihood phylogenetic tree reconstructed using IQTree 3.0.1 showing the relationship between eukaryotic ZDHHC13 and ZDHHC17 orthologue sequences. Line colour indicates bootstrap values as in the legend, active site motif sequences are in brackets, and species names are coloured as follows: vertebrates in black, cyclostomes in blue, insects in maroon, protozoans in grey, nematodes in orange, fungi in brown, and plants in green. Two dashes indicate where branch lengths ≥ 4 were shortened by half for fit.

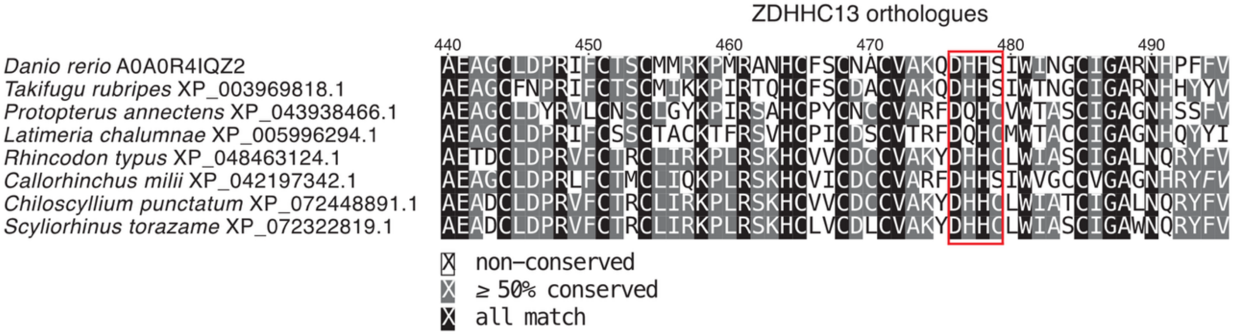


**Figure S3**. Multiple sequence alignment of the cysteine-rich domain of ZDHHC13 fish orthologues with the catalytic motif highlighted with a red box. The alphanumeric code for *Danio rerio* is the UniProtKB accession number and the rest are RefSeq/NCBI Protein accession numbers.

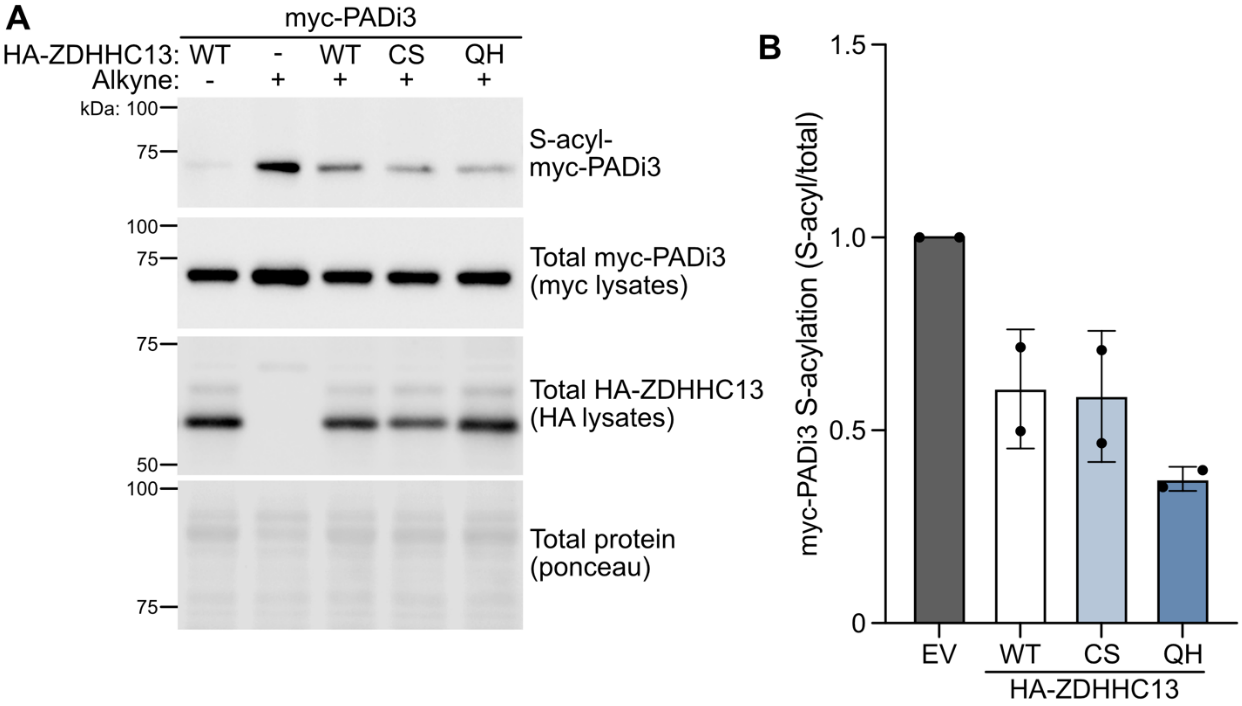


**Figure S4.** ZDHHC13 does not S-acylate PADi3. (A) HEK293T cells expressing myc-PADi3 alone (EV or -) or with HA-ZDHHC13 WT, CS, or QH were labelled with alkyne-palmitate or palmitate (negative control) for one hour. S-acylated proteins were purified from lysates using click chemistry and detected by immunoblot using a myc antibody (top panel). Total myc and HA levels in parent lysates were also determined (middle panels) as well as total protein using ponceau stain (bottom panel). (B) Quantified data from A, showing S-acyl/total levels of myc-PADi3 normalized to EV (N=2).

**Table 1.** Key Resources table.

| DNA | | |
|---|---|---|
| **Plasmid** | **Source** | **Species & promoter** |
| pFEW-HA-*Zdhhc13* | Dr. Gareth Thomas, Temple University[1] | Mouse cDNA; EF1α (human eukaryotic elongation factor 1a) |
| pFEW-HA-*Zdhhc17* | Dr. Gareth Thomas[1] | Mouse cDNA; EF1α |
| pFEW-HA-*Zdhhc20* | Dr. Gareth Thomas[1] | Mouse cDNA; EF1α |
| pFEW-myc-*PADI3* | cDNA from Dr. Anirban Banerjee, National Institutes of Health; subcloned by PCR into XhoI/NotI sites of myc-pFEW (Dr. Gareth Thomas)[1] | Human cDNA; EF1α |
| pFEW-HA (empty vector) | Dr. Gareth Thomas[1] | EF1α |
| HTT$^{1-588}$-peYFPN1 | Dr. Ray Truant, McMaster University[2] | Human cDNA; cytomegalovirus (CMV) |
| LentiCas9-Blast | Dr. Feng Zhang (Addgene plasmid #52962; http://n2t.net/addgene:52962; RRID: Addgene_52962).[3] | *Streptococcus pyogenes*; EF1α |
| pCLIP-DUAL-SFFV-ZsGreen-sgRNA | University of Ottawa Genome Editing and Molecular Biology (GEM) Facility[4] | Human gene targeting gRNAs; U6 RNA polymerase III |
| **gRNA sequences** | | |
| **Gene** | **Catalogue number & sequences** | |
| *ZDHHC13* | TEDH-1090120; gRNAa CATGTATGCAACTGCTGTG and gRNAb GAGGTGGCTGCAGAAATGCG | |
| *ZDHHC17* | TEDH-1090140; gRNAa AATTCTGGATGTATGTGACG and gRNAb GCTATTGTGGATCAACTTGG | |
| *eGFP* | Non-targeting negative control, gRNAa CGAGGAGCTGTTCACCGGGG and gRNAb CAACTACAAGACCCGCGCCG | |
| **Media, reagents, & chemicals** | | |
| **Reagent** | **Catalogue number** | **Manufacturer** |
| Q5 site-directed mutagenesis kit | #E05525 | New England Biolabs (NEB), Ipswich, MA, USA |
| DMEM | #319-015-CL | Wisent Inc., Saint-Jean-Baptiste, QC, Canada |
| Penicillin-Streptomycin | #450-201-EL | Wisent Inc. |
| L-glutamine | #609-065-EL | Wisent Inc. |
| Puromycin | #PUR333.25 | BioShop Canada Inc, Burlington, ON, Canada |
| Fetal bovine serum | #12483020 | Gibco, Waltham, MA, USA |
| Charcoal stripped fetal bovine serum | #F6765 | MilliporeSigma |
| Alkynyl-palmitate (15-hexadecynoic acid) | #CCT-1165 | Vector Laboratories, Newark, CA, USA |
| Palmitate | #P5585 | MilliporeSigma |
| Fatty acid free BSA | #A7030 | MilliporeSigma |
| Roche cOmplete EDTA-free protease inhibitor cocktail | #11873580001 | MilliporeSigma |
| *DC* protein assay | #5000111 | Bio-Rad Laboratories Inc, Hercules, CA, USA |
| Iodoacetamide | #122270050 | ThermoFisher Scientific, Waltham, MA, USA |
| Biotin azide plus | #CCT-1488 | Vector Laboratories |

| | | |
|---|---|---|
| TBTA | #678937 | MilliporeSigma |
| $CuSO_4$ | #209198 | MilliporeSigma |
| Sodium L-ascorbate | #7631 | MilliporeSigma |
| Pierce high capacity NeutrAvidin agarose beads | #PI29204 | ThermoFisher Scientific |
| Hydroxylamine hydrochloride | #159417 | MilliporeSigma |
| MMTS | #64306 | MilliporeSigma |
| Biotin-HPDP | #A8008 | APExBIO Technology LLC, Houston, TX, USA |
| SYPRO Ruby Protein Blot Stain | #S11791 | ThermoFisher Scientific |
| Precision Plus Protein Dual Color Standards | #1610394 | Bio-Rad |
| High molecular weight marker (blue native PAGE) | #17044501 | Cytiva, Marlborough, MA, USA |
| Clarity Western ECL substrate | #1705061 | Bio-Rad |
| SuperSignal West Femto Maximum Sensitivity Substrate | #PI-34096 | ThermoFisher Scientific |
| 4-15% Mini-PROTEAN TGX Stain-Free Protein Gels | #4568086 | Bio-Rad |
| Coomassie G-250 | #CBB555 | Bioshop Canada Inc |
| **Antibodies** | | |
| **Target, type, & dilution** | **Catalogue number** | **Manufacturer** |
| Anti-HA rabbit monoclonal; 1:2000 | #S3724 | Cell Signaling Technologies, Danvers, MA, USA |
| Anti-myc mouse monoclonal; 1:1000 | #2276 | Cell Signaling Technologies |
| Anti-GFP rabbit polyclonal; 1:5000 | #EU1 | Eusera, Edmonton, AB, Canada |
| Anti-ZDHHC13 rabbit polyclonal; 1:500 | #24759-1-AP | Proteintech Group, Inc, Rosemont, IL, USA |
| Anti-rabbit polyclonal anti-ZDHHC17; 1:500 | #H7414 | MilliporeSigma |
| HRP-conjugated anti-rabbit; 1:5000 | #AP182P | MilliporeSigma |
| HRP-conjugated anti-mouse; 1:5000 | #7076 | Cell Signaling Technologies |

| **Buffers** | |
|---|---|
| **Buffer** | **Composition** |
| 2x HBS | 270 mM NaCl, 1.5 mM $Na_2HPO_4 \cdot 7H_2O$, 40 mM HEPES, 10 mM KCl, 10 mM D-glucose, pH 7.0 |
| RIPA buffer | 50 mM HEPES (4-(2-hydroxyethyl)-1-piperazineethanesulfonic acid) pH 7.4, 0.5% sodium deoxycholate, 150 mM NaCl, 1% Igepal CA-630, 0.1% SDS (sodium dodecyl sulfate), 2 mM $MgCl_2$, 1mM iodoacetamide |
| 2% SDS buffer | 50 mM HEPES pH 7.0, 2% SDS, 1 mM EDTA |
| Dilution buffer | 50 mM HEPES pH 7.0, 1% Triton X-100, 1 mM EDTA, 1 mM EGTA [ethylene glycol-bis(β-aminoethyl ether)-N,N,N′,N′-tetraacetic acid], 1 μg/mL leupeptin and 1 mM benzamidine |
| Hydroxylamine elution buffer | 1M $NH_2OH$, 50mM HEPES pH 7.4, 150mM NaCl, 0.1% SDS |
| 4% SDS buffer | 50 mM Tris [tris(hydroxymethyl)aminomethane] pH 7.5, 4% SDS, 5 mM EDTA |

| | |
|---|---|
| Elution buffer | 150 mM NaCl and 0.2% SDS in dilution buffer |
| Hypotonic buffer | 10 mM HEPES pH 7.4 |
| Homogenization buffer | 250 mM HEPES pH 7.4, 750 mM NaCl, 25 mM $MgCl_2$, 2.5 mM dichloro-diphenyl-trichloroethane |
| Blue-native page lysis buffer | 10 mM Tris pH 7.0, 500 mM ε-aminocaproic acid, 20 mM NaCl, 2 mM EDTA, 10% glycerol, 1% N-dodecyl-β-D-maltoside |
| Cathode buffer | 25 mM Tris, 0.1924 M glycine, 0.005% Coomassie G-250 |
| Anode buffer | 25 mM Tris, 0.1924 M glycine |
